# Dependence of an aquatic insect population on contemporaneous primary production

**DOI:** 10.1101/2021.03.22.436388

**Authors:** Joseph S. Phillips, Amanda R. McCormick, Jamieson C. Botsch, Anthony R. Ives

## Abstract

1. Characterizing the dynamics of energy flow through ecosystems requires quantifying the degree to which primary and secondary production are coupled. This coupling is expected to be tight in ecosystems with high internal production relative to external carbon and energy inputs.
2. We experimentally quantified the dependence of aquatic insect emergence on fresh primary production, specifically for the midge population in Lake Mývatn, Iceland. Using field mesocosms, we manipulated algal primary production by reducing light availability via shading. We then used dissolved oxygen incubations to estimate fluxes of carbon through photosynthesis (i.e., gross primary production or “GPP”) over the course of the experiment.
3. We found that elevated GPP was associated with higher emergence rates of adults, as judged both by comparison of emergence across the experimental shading treatments and estimates of in situ GPP within the mesocosms. Furthermore, larger adults emerged earlier than smaller ones, suggesting that asymmetries in resource availability among individuals affected the timing of emergence. Nonetheless, midge emergence was substantial under light-limiting conditions, indicating that while midges benefit from primary production contemporaneous with larval development, they are also capable of completing their life cycles on carbon already existing in the organic matter pool.
4. Our results show that even in systems with limited allochthonous inputs, contemporaneous primary production may be not necessary for high secondary production and insect emergence. Instead, consumers can develop from consumption of biomass derived from past autochthonous primary production. This suggests that primary production and consumer dynamics can be partially decoupled in time in systems that depend on internal production.

## Introduction

The transfer of carbon and energy across trophic levels plays a central role in governing population, community, and ecosystem dynamics (Hairston Jr and Hairston Sr, 1993; Carpenter et al., 2001; Chapin et al., 2006). In freshwater ecosystems, much attention has been devoted to tracing pathways of energy flow (Lindeman, 1942; Cole et al., 2002; Schmidt et al., 2007), often emphasizing the relative contribution of external (allochthonous) versus internal (autochthonous) carbon and energy inputs (Kritzberg et al., 2004; Brett et al., 2009; Rosi-Marshall et al., 2016).

The allochthonous versus autochthonous dichotomy is paralleled by the difference between “fresh” or contemporaneous primary production, whereby secondary production is fueled by consumption of recently assimilated carbon, versus “stored carbon” that is the result of past primary production (Wetzel, 1995; Venarsky et al., 2014). Stored carbon can include living tissue from past production, detritus, or more recalcitrant material, although the latter is unlikely to contribute directly to secondary production. Although detritus has long been recognized as an important resource for consumers (Lindeman, 1942; Wetzel, 1995), relatively few studies have assessed the extent to which secondary production relies on contemporaneous primary production (but see Solomon et al., 2008; Huryn et al., 2014).

The external vs. internal and the fresh vs. stored dichotomies play a central role in determining the extent to which primary and secondary production are dynamically coupled within a given ecosystem. Primary consumers are expected to rely more heavily on allochthonous inputs when allochthonous inputs are high relative to autochthonous production (Wallace et al., 1997; Marczak et al., 2007; Rosi-Marshall et al., 2016). Alternatively, when autochthonous production is high, consumers are expected to depend more heavily on autochthonous production as it is often makes for higher quality food (Marcarelli et al., 2011). When both contemporary autotrophic production and allochthonous inputs are low, consumers either need to temporarily slow their metabolic rates (e.g., through diapause) or rely more heavily on the carbon stored in the system from past autochthonous production (Bellamy and Bauer, 2017). For example, spatial patterns of resource utilization by benthic macroinvertebrates in a temperate lake show that reductions in light and benthic primary production associated with depth are paralleled by a shift in resource utilization from autotrophic production to autochthonous detritus (Solomon et al., 2008).

In this study, we experimentally quantified the dependence of aquatic insect emergence on fresh primary production, using the midge *Tanytarsus gracilentus* (Diptera: Chironomidae) in Lake Mývatn, Iceland. This population of *T. gracilentus* fluctuates over several orders-of-magnitude with crashes occurring every 4 to 10 years (Einarsson et al., 2002; Ives et al., 2008). These crashes appear to be proximately driven by food limitation (Einarsson et al., 2002), and the population fluctuations are associated with variation in diatom pigments (Einarsson et al., 2016) and stable carbon isotopes (McCormick et al., In Review) in a manner consistent with consumer-resource dynamics. Moreover, laboratory experiments have shown that growth and survival of *T. gracilentus* larvae are promoted under conditions conducive to algal production (Wetzel et al., 2021). However, the midge larvae not only feed on living algal cells but also on bacteria and detritus (Ingvason et al., 2002, 2004), potentially decoupling their dynamics from fresh primary production. Indeed, a mathematical model of their population dynamics suggests that detritus could introduce time-lags that facilitate the depletion of algal biomass, which in turn may set the stage for their high-amplitude and aperiodic fluctuations (Ives et al., 2008).

The Mývatn ecosystem experiences high autochthonous production relative to allochthonous inputs (Einarsson et al., 2004). However, it also shows dramatic intra- and interannual variation in benthic primary production, which is the primary source of autotrophic production available to *T. gracilentus*. A primary driver of this variability is shading by cyanobacteria blooms, which occur in the main basin irregularly (Einarsson et al., 2004; Phillips, 2020). During these blooms, cyanobacteria densities can eliminate all light reaching the benthos, such that benthic production is drastically reduced (McCormick et al., 2021). The benthos at Mývatn consists of a thick layer of unconsolidated diatomaceous ooze, which may serve as an important resource for *T. gracilentus* during these blooms, although it is also possible that they feed on settling carbon from the blooms.

Using field mesocosms, we experimentally manipulated algal primary production by reducing light availability via shading, and we then used dissolved oxygen incubations to estimate fluxes of carbon through photosynthesis over the course of the experiment. Because consumers can strongly reduce biomass of benthic algae (Hillebrand, 2009), the rate of carbon fixation through primary production rather than estimates of algal biomass is likely to be more relevant in describing the dependence of midges on contemporary autotrophic production (Rüegg et al., 2021). The mesocosms were colonized by first-instar midge larvae, and then they incubated in the lake until the midges began to emerge as adults, enabling us to quantify the response of midges to variation in primary production throughout their life cycle. Our results shed light on the coupling of contemporaneous primary and secondary production, with potential consequences for the dynamics of consumer populations.

## Methods

### Study site

Mývatn is a large shallow lake located in northeast Iceland (65°40’N 17°00’W), with a surface area of 37km^2^ and a mean depth in the main basin of ≈2.5m. (Einarsson et al., 2004). The lake is fed by nutrient-rich springs, with inputs of N, P, and Si = 1.4, 1.5, and 340 g m^−2^ year^−1^ (Ólafsson, 1979). Due its shallowness, relatively clear water, and high nutrient inputs, Mývatn supports high benthic primary production, chiefly comprising diatoms of the family Fragilariaceae (Ólafsson, 1979; McCormick et al., 2019, 2021).

Mývatn’s high benthic production supports large populations of midges (Chironomidae), with the sediment-dwelling midge *T. gracilentus* being the dominant benthic/epibenthic invertebrate (Lindegaard and Jónasson, 1979). *T. gracilentus* has four larval instars, and the second through fourth instars build silk tubes in the lake sediment where they feed on algae and detritus (Ingvason et al., 2002, 2004). In most years the population is clearly bivoltine (Lindegaard and Jónasson, 1979). Adults from the first generation emerge in early June and their offspring comprise the second generation that emerges in late July. The offspring of the second generation overwinter as third-instar larvae until the next June emergence. *T. gracilentus* belongs to the tribe Tanytarsini, and in this study larval midges were only identified to this resolution. However, the vast majority of the Tanytarsini from the areas of Mývatn relevant to this study are likely *T. gracilentus*.

### Study design

To quantify the dependence of *T. gracilentus* emergence on sediment variation and contemporaneous primary production, we conducted a field mesocosm experiment. The experiment was originally conceived with a 2 × 3 × 2 design, with two shading treatments (shaded vs. light) to produce variation in primary production, three sediment sources (sites E1, E3, and E4), and two midge abundance treatments (exclusion and no exclusion). We used a total of 60 mesocosms evenly distributed across the treatments.

The midge exclusion was intended to quantify the effect of midges on benthic primary production, which previous studies at Mývatn have shown to be large and generally positive (Herren et al., 2017; Phillips et al., 2019). However, the exclusion treatment failed to exclude midges (see below). Because including the midge exclusion treatment did not substantively affect the statistical results, for clarity we present analyses without the exclusion treatment in the main text, although we include analogous analyses with the exclusion treatment in the *Supplementary Materials* (Tables S2-S5). Therefore, the experiment effectively had a 2 × 3 design.

The mesocosms were made from clear acrylic tubes (33cm height x 5cm diameter) sealed from the bottom with foam stoppers and filled with sediment collected from Mývatn. On 6 June 2016, we collected sediment with Kajak corers from the three sites (E1, E3, E4; labeled west to east) spanning the main basin (locations shown in Figure S1). These sites were selected because they had a wide range of Tanytarsini larval densities assessed during a survey on 29-31 May 2016. Specifically, E4 had the highest Tanytarsini density (mean ± standard error of 23, 428 ± 7, 713 m^−2^), followed by E1 (5, 432 ± 612) and then E3 (2, 241 ± 831). We expected larval density to be associated with sediment quality for larval development in the experiment for two reasons. First, midge abundance could reflect the quality of the sediments for midge growth and survival at a site. Second, the abundance of midges could affect the sediment in a way that influences development and survival in the next generation (i.e., larvae colonizing the experimental mesocosms; see below).

We separated the top 5cm and next 10cm of each core, and then combined sediment from multiple cores for each site × sediment layer combination. Next, we sieved the top layer sediment through 125 µm mesh and the bottom layer sediment through 500 µm to remove sediment-dwelling midges; different mesh sizes were used because the majority of midges (particularly those smaller than 500 µm) reside in the top layers of sediment. Following sieving, we allowed the sediment to settle for three days in cool and dark conditions before stocking the mesocosms on 9 June 2016. For each site, 20 mesocosms were each filled with 10cm of bottom-layer sediment, followed by 5cm of top-layer sediment from the appropriate site. The actual depth of the sediment layer in the mesocosms varied within a 5cm range, depending on the extent to which the sediment settled; the resulting variation in the water column depth was accounted for in the analyses. Finally, we wrapped the bottom portion of each mesocosm corresponding to the sediment layer with 4 layers of black plastic to exclude lateral light from reaching the sediment.

On 9 June 2016, we haphazardly distributed the mesocosms across 5 experimental racks and deployed them in a sheltered bay on the south side of Mývatn approximately 150m from shore at a depth of 1.7m. To deploy the mesocosms, we added water to the tops of the mesocosms collected from the deployment site. This resulted in modest resuspension of sediment within the mesocosms; therefore, we allowed the sediment to settle for several minutes before deployment. Half of the mesocosms were left with open tops to allow colonization from *T. gracilentus* eggs and larvae resulting from mating swarms of *T. gracilentus* that were active at the time. The tops of the remaining mesocosms were covered with 63 µm mesh to prevent colonization; these exclusions failed, both because some of the mesh covers detached from the mesocosms and because water used to fill the mesocosms likely contained eggs and first instar larvae.

We left the mesocosms in the lake for 15 days to allow colonization by *T. gracilentus* and equilibration of the mesocosms with the ambient environment. Then, on 26 June 2016 we measured gross primary production (GPP), ecosystem respiration (ER), and net ecosystem production (NEP), which are collectively known as “ecosystem metabolism” (see details in *Methods : Metabolism measurements*). We then established the “shaded” treatment by wrapping 4 layers of black plastic around the water columns of half of the mesocosms. The shading only blocked light from the sides, so the shaded treatment allowed some light to reach the sediment (around 2% of the ambient light available to the light mesocosms). Finally, we removed the mesh covers from the midge-exclusion tubes before redeploying the mesocosms on the lake bottom.

After incubating in the lake for an additional 16 days, on 12 July 2016 we repeated the ecosystem metabolism measurements. At this time, we expected that the *T. gracilentus* that colonized the mesocosms were nearing emergence. Therefore, on 12 July we attached a mesh cage to the top of each mesocosm to capture emerging adults. Over the next 14 days, we stored the mesocosms outdoors in tents made of mosquito mesh to reduce light and minimize external contamination by adult midges. To moderate temperature in the mesocosms, we kept them in water baths (routinely refilled with cold tap water) deep enough to submerge the sediment and most of the water column, while leaving the tops exposed so that pupae could emerge as adults. We collected any midges that emerged from the mesocosms approximately daily and identified the male adults to species (females are difficult to identify). The large majority of these midges were *T. gracilentus*, particularly after the first couple of days of collection. We then measured the length of a single wing from each male *T. gracilentus*, measuring from the arculus to the wingtip as in Einarsson et al. (2002). After 14 days, the emergence rates began to attenuate (Figure S3), at which point we terminated the experiment by sieving the sediment from each mesocosm through 125 µm mesh and picking out any remaining midge larvae. We measured the head capsule widths of the Tanytarsini larvae, which we used to classify individuals into discrete instars (Figure S2).

### Metabolism measurements

On 26 June and 12 July 2016 (days 17 and 33 of the experiment), we quantified ecosystem metabolism by measuring the change in dissolved oxygen in the water column of each mesocosm during sequential incubations under light and dark conditions, following Phillips et al. (2019). These estimates take advantage of the fact that GPP releases oxygen while ER consumes it, and fluxes of oxygen for both of these processes correspond with approximately equal fluxes of carbon (Hall and Hotchkiss, 2017). All mesocosms, regardless of their experimental shading treatment, were incubated under both dark and light conditions for the purpose of separating GPP and ER. For the light incubations, we measured the initial dissolved oxygen concentration (DO; mg L^−1^) of the water column of each mesocosm with a handheld probe (ProODO, YSI, Yellow Springs, Ohio, USA), gently stirring to homogenize the water. We then sealed the mesocosms with rubber stoppers and deployed them on the lake bottom (1.7m) for the duration of the incubation lasting between 4 and 6 hours. After the incubation period, we measured the final DO. We repeated this procedure (using the final reading from the first incubation as the initial reading for the second), but with 4 layers of black plastic wrapped around the water column of each mesocosm to eliminate light. On each sampling day, we measured photosynthetically active radiation (PAR) at multiple depths (differing depending on the sample day) throughout the incubation period using a Li-192 Quantum Underwater Sensor (Li-COR, NE, USA). We used these measurements to estimate the light level at the incubation depth (1.7m) during the incubation period for each experimental rack by assuming an exponential attenuation of light with depth.

For each incubation, we calculated the areal flux rate of DO across a cross section of the mesocosm by multiplying the change in DO concentration by the water column depth within the mesocosm and dividing by the incubation duration; this yielded measurements in units of g O_2_ m^−2^ h^−1^. Because metabolism in the mesocosm water columns was likely negligible relative to that of the sediment (Phillips et al., 2019), this can be interpreted as the flux of oxygen across the sediment surface. We assumed that ER remained approximately constant during the two incubations in light and dark conditions for each mesocosm on a given date. Therefore, we interpreted the DO fluxes in the light incubations as NEP, fluxes in the dark as −|ER|, and calculated GPP as NEP + ER.

Our estimates of GPP were based on NEP measurements under ambient light conditions and without shading (regardless of the shading treatment). However, we were also interested in the GPP that occurred under the light conditions experienced by the shaded mesocosms during the main experiment, which were substantially lower than the ambient light levels but not completely dark as the mesocosms were only shaded from the sides (when not stoppered as during the metabolism incubations). Therefore, we extrapolated GPP under the lower in situ light levels (*GPP_insitu_*) using a photosynthesis-irradiance curve of Michaelis-Menten form:

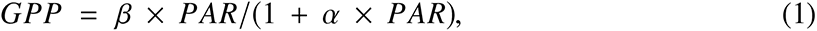

where PAR is the light level, β is the maximum rate of increase of GPP with increasing light and α is the rate at which the curve saturates (Jassby and Platt, 1976). We set α equal to the value reported in Phillips et al. (2019), which was based on a separate experiment conducted at Mývatn that used the same mesocosm setup as the present study but with an experimentally generated gradient of light levels. By using this value, we were able to calculate β for the present study as

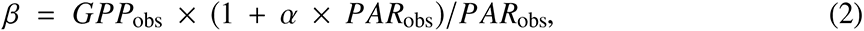

with *GPP*_obs_ and *PAR*_obs_ corresponding to values observed during the metabolism incubations on a given experimental day (17 or 33). Finally, we estimated *GPP*_in situ_ under ambient light conditions as

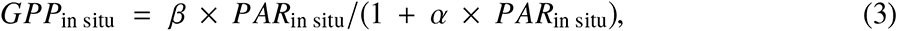

where *PAR*_in situ_ was calculated from the observed PAR times the fractional reduction in light due to the shading treatment for the corresponding mesocosm. We performed these calculations for both days on which GPP was measured, which we averaged to yield a single estimate of in *GPP*_in situ_ for each mesocosm.

### Data analysis

We analyzed variation in GPP across the experimental treatments with a linear mixed model (LMM). We included fixed effects for sediment source site (3 levels), shading treatment (2 levels), incubation date (2 levels), and their two-way interactions. We also included a fixed main effect of PAR during the light incubations for each rack-date combination. Random intercepts grouped by experimental rack (5 levels) were included to account for blocking of the measurements. For the analyses presented in the main text, we omitted the “midge exclusion” treatment due to the treatment’s failure to exclude midges. However, in the *Supplementary materials* (Tables S2-S5) we report results of analyses with the exclusion treatment added; the substantive statistical conclusions remain the same regardless.

We analyzed variation in the abundance of Tanytarsini larvae in different instars at the end of the experiment using a single LMM with log(count + 1) as the response variable and fixed effects for sediment source site (3 levels), shading treatment (2 levels), instar (4 levels), and their two-way interactions. We also included the three-way interaction between source site, shading, and instar to allow each instar to respond separately to the interaction between the experimental treatments. Random intercepts grouped by experimental rack (5 levels) and mesocosm (60 levels) were included to account for blocking and the multiple measurements for each mesocosm. Despite the fact that the different instars likely represented different generations, we fit them in a single model because the exact delineation between the two generations (offspring from either spring and summer emergences) was unclear and because the data were collected from a single sampling process. However, we complemented this analysis by fitting LMMs to each instar separately, including fixed effects for sediment source site (3 levels), shading treatment (2 levels), and their two-way interactions

To determine the effect of experimental treatments on total emergence of *T. gracilentus* over the 14-day sampling period, we fit an LMM for log(total count + 1) from each mesocosm as the response variable, with main effects and two-way interaction for sediment source site (3 levels) and shading treatment (2 levels). Random intercepts grouped by experimental rack (5 levels) were included to account for blocking of the measurements. Finally, we analyzed variation in wing length by fitting an LMM with wing length as the response variable, main effects for sediment source site (3 levels), shading treatment (2 levels), emergence day (continuous), and their two-way interactions. Random intercepts grouped by experimental rack (5 levels) and mesocosm (60 levels) were included to account for blocking and the multiple measurements for each mesocosm.

Analyses were performed in R 4.0.3 (R Core Team, 2020). We fit the LMMs using the lme4 package (Bates et al., 2015). *P*-values for the LMMs were calculated from *F*-tests with the Kenward-Roger correction, using the car package (Fox and Weisberg, 2019). For each model we report *P*-values from both Type-III and Type-II tests. Type-III tests evaluate the main effects from a full model with interactions included, which is useful because dropping terms from a model can lead to inflated coefficient estimates and Type-I errors. However, an overparameterized model can also yield misleading inferences due to the increased noisiness of the parameter estimates. Therefore, it is also useful to consider Type-II tests that drop all interactions before assessing the significance of the main effects. To characterize the overall effects of the categorical predictors, we report the estimated marginal mean response for each predictor level averaged across the other predictors and interactions in the model matrix, using the emmeans package (Lenth, 2020). These means were calculated on the transformed scale when relevant, and therefore were on the same scale as the coefficient estimates. For interactions and continuous predictors, we report coefficients as effects relative to the model intercept. For the LMMs, PAR was centered on its means and divided by two standard deviations to make its corresponding effect size estimates comparable to those of the categorical predictors. All predicted means and coefficients were estimated from the full models including interactions and are reported with ± one standard error based on the model-estimated variance-covariance matrix.

## Results

The main goal of this study was to quantify the effects of benthic primary productivity on secondary production by *T. gracilentus*, measured from growth and survival. Our mesocosm experiment had two treatments: shading to vary the magnitude of in situ gross primary production (GPP), and sediment source (E1, E3, or E4) representing variation in naturally occurring midge abundances. Variation in midge abundance could reflect differences in sediment quality, could create differences in sediment quality, or both.

GPP measured under ambient light (i.e., with the experimental shades removed), did not differ between the shading treatments (shaded: 0.14 ± 0.01; light: 0.14 ± 0.01) (Figure 1; Table 1). No shading treatment difference was expected for the first incubation, because the treatment was not applied until after the incubation (Day 17). However, the interaction between time and shading treatment (0.0007 ± 0.009) was also non-significant. These results suggest that shading did not reduce the potential for photosynthesis by benthic primary producers. GPP varied across sediment sources according to the Type II test (Table 1), with GPP being highest with sediment from E3 (0.16 ± 0.01), followed by E1 (0.14 ± 0.01) and then E4 (0.12 ± 0.01) (Figure 1). This is opposite of the ordering of larval densities at the three sites from which the sediment was collected at the start of the experiment. While the main effect of sediment source was not statistically significant according to the Type III test, there was a marginal interaction between sediment source and incubation date with GPP from mesocosms with E4 sediment increasing less than the other two sites between days 17 and 33. Together, these results suggest that there were differences in the photosynthetic capacity of the sediment from different sites that persisted through time despite the influence of in situ conditions. While GPP under ambient light did not vary by shading treatments, the average in situ light levels in the shaded treatment (1.71 *µ*mol-photons m^−2^ s^−1^) were much lower than in the light treatment (87.2 *µ*mol-photons m^−2^ s^−1^), such that the estimated *GPP_insitu_* was ≈20 times higher in the light versus the shaded treatment.

**Figure 1:**
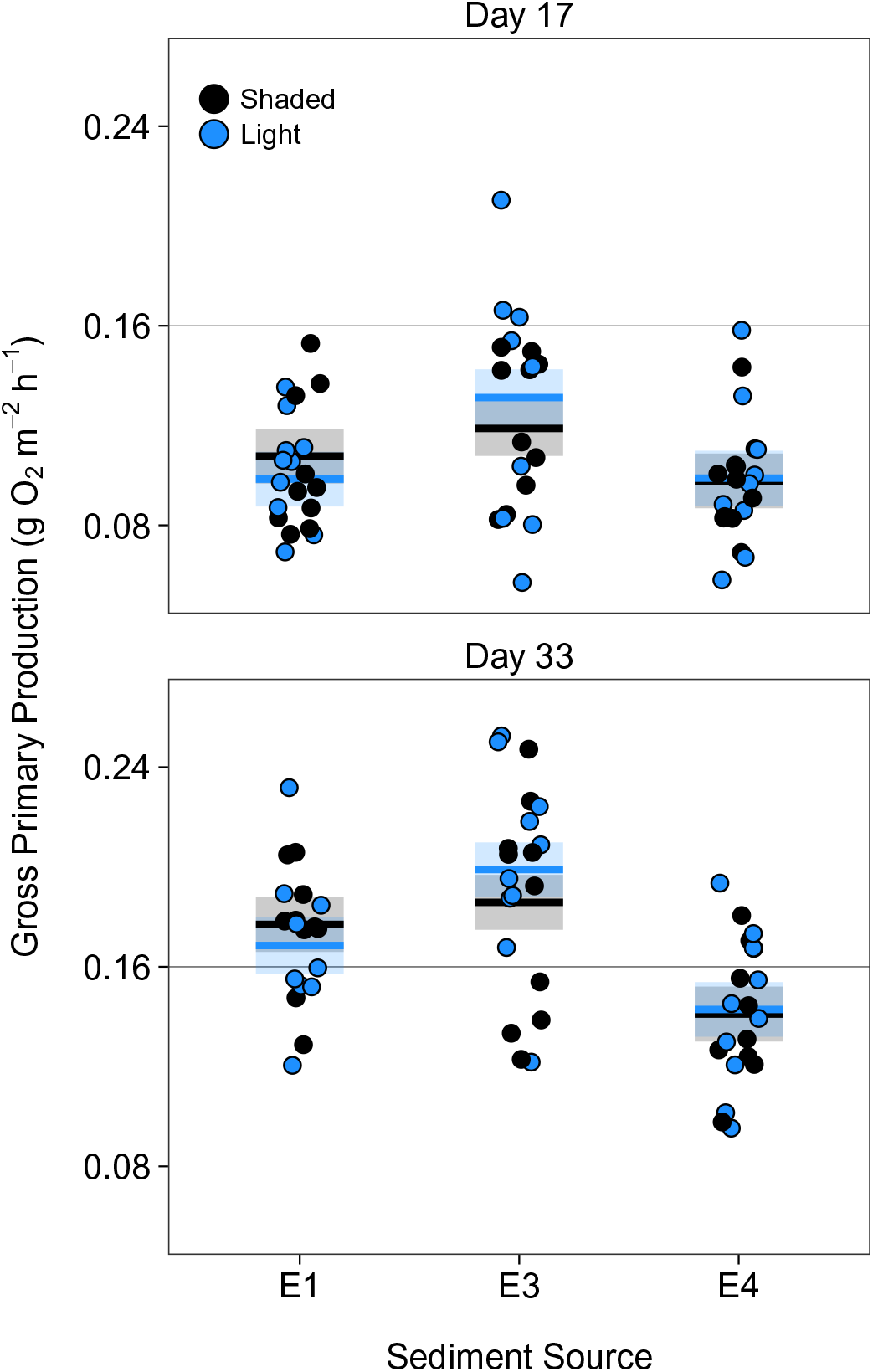
GPP from the mesocosm experiment. Points show the data, standardized to the mean PAR level using the coefficient estimate from the LMM. The lines show means from the full LMM (including all interaction terms), and the shaded regions show standard errors.

**Table 1:** ANOVA for mesocosm GPP. *P*-values are from *F*-tests with the Kenward-Roger correction.

| Term | Type III |  | Type II |  |
| --- | --- | --- | --- | --- |
| | $F_{ndf,ddf}$ | $P$ | $F_{ndf,ddf}$ | $P$ |
| PAR | 19<sub>1,56.6</sub> | <0.001 | 19<sub>1,56.6</sub> | <0.001 |
| site | 1.5<sub>2,74</sub> | 0.229 | 13<sub>2,49.7</sub> | <0.001 |
| shading | 0.61<sub>1,66.8</sub> | 0.439 | 0.094<sub>1,50.5</sub> | 0.760 |
| date | 37<sub>1,54.9</sub> | <0.001 | 52<sub>1,56.7</sub> | <0.001 |
| site × shading | 0.98<sub>2,50.1</sub> | 0.383 | | |
| site × date | 3.4<sub>2,54</sub> | 0.040 | | |
| shading × date | 0.0067<sub>1,54</sub> | 0.935 | | |

At the end of the experiment, larval abundance varied across instar, with second and third instars being most abundant (Figure 2; Table 2). However, neither sediment source nor shading treatment had statistically significant main effects or interactions, including the two- and three-way interactions with instar. When separate models were fit to each instar separately, the Type II test showed marginally negative effects of shading on the abundance of third and fourth instars (Table S1). Together, the results suggest that while it is possible that reduced light levels from shading resulted in fewer third and fourth instars, the magnitude of these effects was low and uncertain.

**Figure 2:**
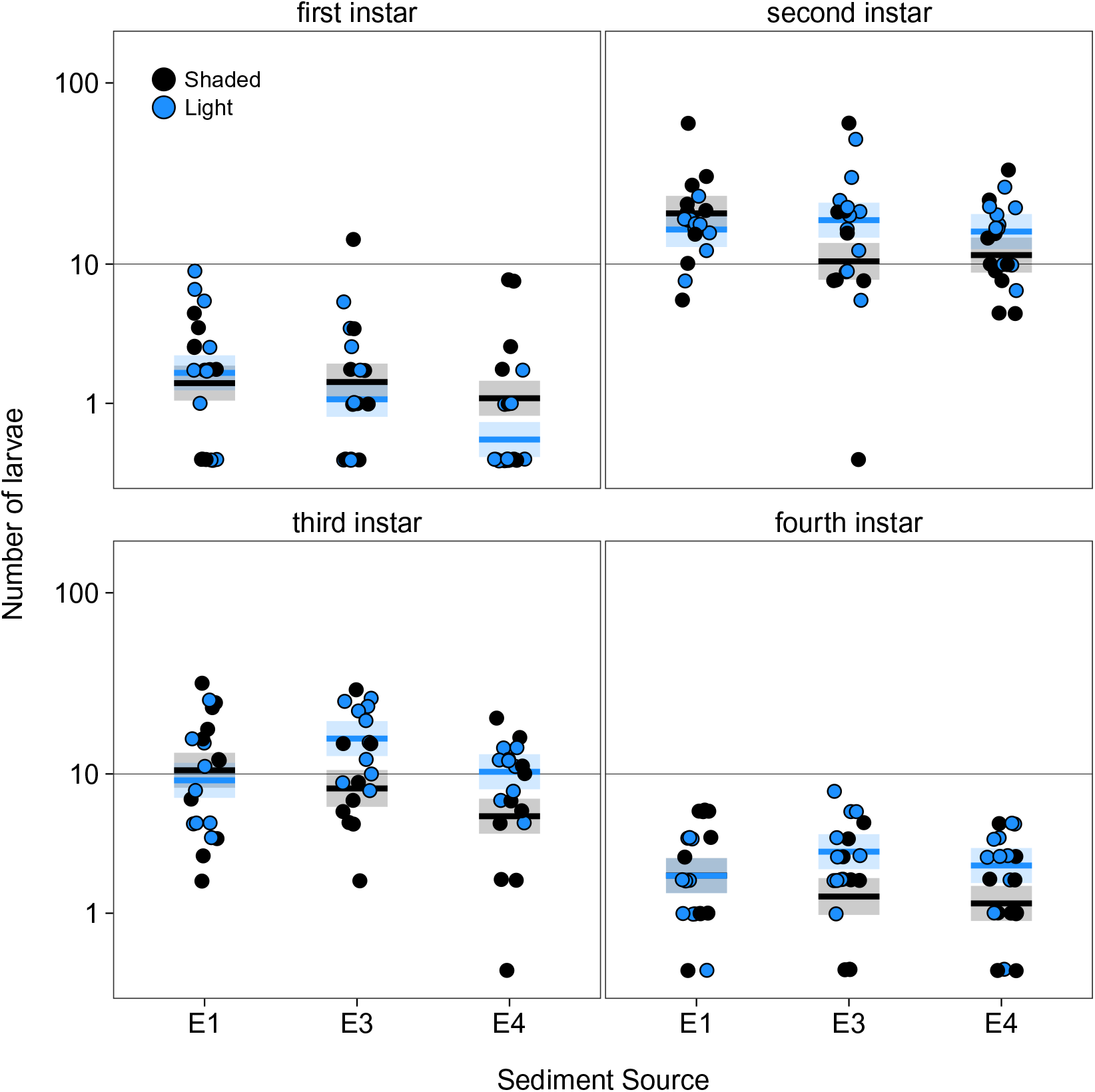
Abundance of Tanytarsini larvae from the mesocosm experiment. Points show the data, lines show means from the full LMM (including all interaction terms), and the shaded regions show standard errors.

**Table 2:** ANOVA for abundance of Tanytarsini larvae in the mesocosms. Counts were log-transformed, with 1 added to all values to accommodate zeros. *P*-values are from *F*-tests with the Kenward-Roger correction.

| Term | Type III |  | Type II |  |
| --- | --- | --- | --- | --- |
| | $F_{ndf,ddf}$ | $P$ | $F_{ndf,ddf}$ | $P$ |
| site | 0.28<sub>2,186</sub> | 0.758 | 2.6<sub>2,49.1</sub> | 0.086 |
| shading | 0.18<sub>1,188</sub> | 0.672 | 2.4<sub>1,51.4</sub> | 0.126 |
| instar | 28<sub>3,159</sub> | <0.001 | 150<sub>3,159</sub> | <0.001 |
| site × shading | 1.1<sub>2,187</sub> | 0.322 | | |
| site × instar | 0.54<sub>6,159</sub> | 0.775 | | |
| shading × instar | 0.28<sub>3,159</sub> | 0.839 | | |
| site × shading × instar | 1.4<sub>6,159</sub> | 0.204 | | |

Across all experimental treatments, daily emergence of adults began near zero before reaching a peak midway through the sampling period, followed by a decline towards zero by the end (Figure S3). This is consistent with the 14-day sampling period having captured the majority of the potential emerging adults, with the remaining individuals from that generation either dead or remaining as larvae. Total emergence was higher in the light treatment (1.66 ± 0.15, on log-scale) than in the shaded treatment (1.19 ± 0.15) (Figure 3); this effect was statistically significant for the Type II test and marginal for the Type III test (Table 3). Daily and total emergence were highest for sediment from E3 (1.62 ± 0.16), followed by E1 (1.38 ± 0.16) and then E4 (1.28 ± 0.16), the same as the rank-order of GPP from the different sites and the reverse of the larval abundance at the sites in the previous generation. However, these differences were modest and not statistically significant. Overall, these results are consistent with a positive effect of light availability on emergence rates of adult *T. gracilentus*.

**Figure 3:**
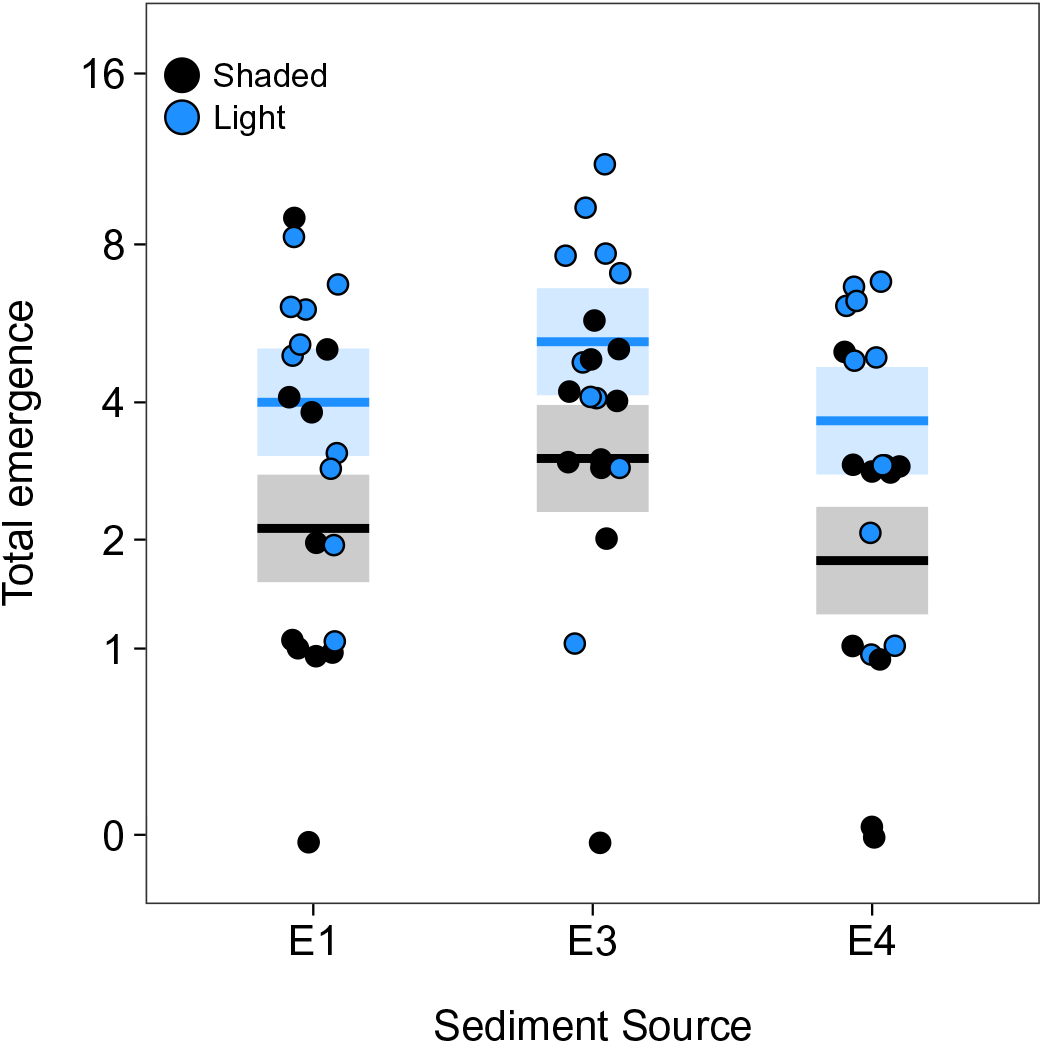
Total emergence of *T. gracilentus* adults from the mesocosm experiment. Points show the data, lines show means from the full LMM (including all interaction terms), and the shaded regions show standard errors.

**Table 3:** ANOVA for number of *T. gracilentus* adults that emerged the mesocosms. Counts were log-transformed, with 1 added to all values to accommodate zeros. *P*-values are from *F*-tests with the Kenward-Roger correction.

| Term | Type III |  | Type II |  |
| --- | --- | --- | --- | --- |
| | $F_{ndf,ddf}$ | $P$ | $F_{ndf,ddf}$ | $P$ |
| site | 1.3<sub>2,50.2</sub> | 0.274 | 2.2<sub>2,50</sub> | 0.127 |
| shading | 3.8<sub>1,51</sub> | 0.057 | 12<sub>1,50.9</sub> | <0.001 |
| site × shading | 0.032<sub>2,50.5</sub> | 0.968 |  |  |

Wing lengths declined with emergence day (−0.08 ± 0.05), and the main effect of emergence day was statistically significant according to the Type II test but not the Type III (Figure 4; Table 4). However, according to the Type III test there was a marginal interaction between emergence day and site, with E3 having a positive interaction with emergence day (0.11 ± 0.05). Taken together, these results indicate that wing lengths generally declined with emergence day, with the possible exception of those from E3 sediment. Beyond the potential interaction between sediment source and day of emergence, wing lengths did not vary substantially by either sediment source or shading treatment.

**Figure 4:**
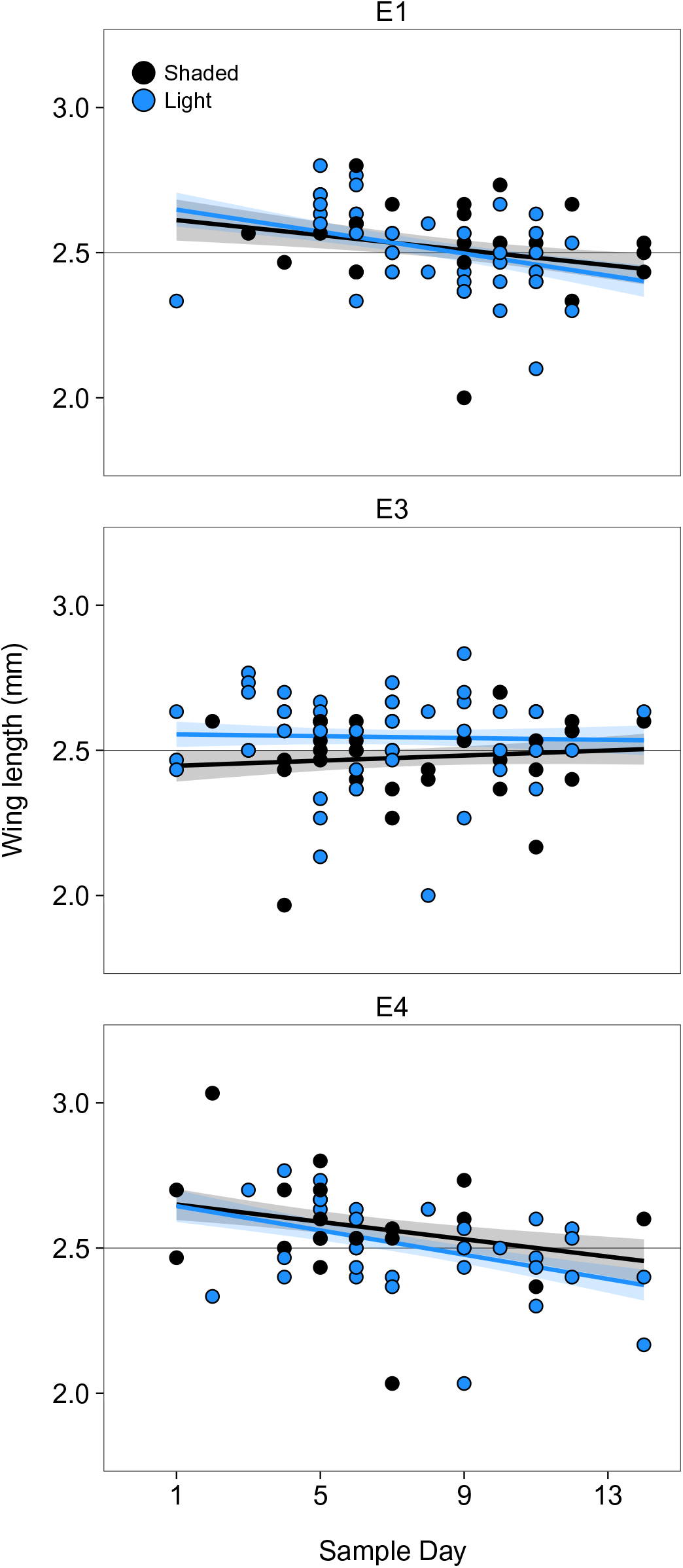
Wing lengths of *T. gracilentus* adults from the mesocosm experiment. Points show the data, lines show means from the full LMM (including all interaction terms), and the shaded regions show standard errors.

**Table 4:** ANOVA for wing lenghts of *T. gracilentus* adults that emerged the mesocosms. *P*-values are from *F*-tests with the Kenward-Roger correction.

| Term | Type III |  | Type II |  |
| --- | --- | --- | --- | --- |
| | $F_{ndf,ddf}$ | $P$ | $F_{ndf,ddf}$ | $P$ |
| date | 2.6<sub>1,201</sub> | 0.109 | 7.2<sub>1,199</sub> | 0.008 |
| site | 1.3<sub>2,59.9</sub> | 0.278 | 0.22<sub>2,37.8</sub> | 0.806 |
| shading | 0.0056<sub>1,49.4</sub> | 0.941 | 0.52<sub>1,43.1</sub> | 0.474 |
| date × site | 3.1<sub>2,198</sub> | 0.047 |  |  |
| date × shading | 0.66<sub>1,197</sub> | 0.420 |  |  |
| site × shading | 1.7<sub>2,46.5</sub> | 0.190 |  |  |

Together, our results suggest that Tanytarsini emergence rates increased with the availability of light, potentially mediated through light effects on in situ GPP. To make this link more explicit, we fit a Poisson generalized linear mixed model (GLMM) with total emergence as the response and estimated in situ GPP as the predictor, which encompassed variation due both to experimental shading and sediment source. We included random intercepts grouped by experiment rack and mesocosm identity to account for blocking and potential overdispersion. The coefficient estimate for the effect of in situ GPP on total emergence was 3.81 ± 1.07 on the log-link scale. This implied that a 0.1 mg O_2_ m^−2^ h^−1^ increase in GPP would result in a 4% increase in the number of emerging midges on the count scale, or a 67% increase in emergence from the light treatment (4.58 individuals per mesocosm) over the shaded treatment (2.75 individuals).

## Discussion

Using field mesocosms experiments, we evaluated the extent to which the *T. gracilentus* of Lake Mývatn depends on fresh primary production during their larval life stages. Elevated GPP was associated with higher emergence rates of adults, as judged both by comparison of emergence across the experimental shading treatments and estimates of in situ GPP within the mesocosms. Furthermore, the fact that larger adults emerged earlier than smaller ones suggests that asymmetries in resource availability among individuals affected the timing of emergence. Nonetheless, *T. gracilentus* emergence was substantial in the shaded treatments, with 2% of the PAR and 8% of the in situ GPP compared to the non-shaded treatments under in situ conditions. This indicates that while *T. gracilentus* benefit from “fresh” primary production, they are also capable of completing their life cycles on carbon already existing in the organic matter pool.

Our finding that light limitation reduced primary production and subsequent emergence is consistent with previous studies (Mayer and Likens, 1987; Lamberti and Resh, 1983; Wellnitz et al., 1996). For example, experimental light reduction in an artificial stream reduced biomass of benthic algae and subsequent growth by snails (Hill and Harvey, 1990; Hill et al., 1995). However, these previous studies used primary producer biomass as an indicator of productivity, in contrast to our study which quantified carbon fixation more directly through fluxes of dissolved oxygen. This is an important distinction, as standing biomass and carbon assimilation rates may not be tightly linked. Indeed, in our experiment there was no difference in GPP between the shaded and un-shaded treatments when both were incubated under ambient light, despite the fact that GPP under in situ conditions was certainly different due to the large difference in ambient light availability. This suggests that either the higher in situ GPP in the un-shaded treatments did not result in net biomass accrual or that some other factor became limiting in the un-shaded treatments. Given the expected reduction in algal biomass due to grazing by midges (Lamberti and Resh, 1983; Wellnitz et al., 1996), it could be that excess algal production under light conditions was consumed by midges, contributing to their higher emergence rates, as seen for other aquatic insects (Mayer and Likens, 1987; Ball and Baker, 1995). It is for this reason that primary productivity per se has been argued to be more relevant than standing biomass for controlling secondary production (Wetzel, 1995; Rüegg et al., 2021).

While emergence rates of adult midges were lower in the shaded treatment, larval abundance was similar between the two treatments. This suggests that food limitation induced by shading primarily manifested in larval growth rates, leading to differences in pupation and emergence. It is also possible that the larvae emigrated at higher rates from the shaded mesocosms; however, if this were induced by poor feeding conditions, it would still be indicative of the importance of fresh primary production. Our result is consistent with previous studies showing that food limitation during the larval stage affects survival and emergence timing of adult Diptera (Collins, 1980; Levi et al., 2014) and Lepidoptera (Kelly and Debinski, 1999). Long-term records of adult *T. gracilentus* at Mývatn show that declining wing lengths preceded population crashes, suggesting that the crashes are driven by food limitation (Einarsson et al., 2002). In our experiment, we did not observed variation in wing lengths between the shading treatments. Given the higher emergence rates from the light treatment, this suggests that increased food availability and consumption was primarily allocated to speed development through the life cycle, rather than increasing body size. Nonetheless, we did observe that larger individuals tended to emerge earlier than smaller ones, which is consistent with asymmetric distribution of food resources affecting the timing of emergence (Collins, 1980).

In our experiment, primary production differed among sediment source locations, and this variation was mirrored in emergence rates. However, the rank-order of variation in primary production was opposite that of ambient densities of Tanytarsini larvae at the sediment source locations when the sediment was collected. Previous work in both Mývatn and other systems has shown that tube-building midge larvae can stimulate benthic primary production by providing a 3-dimensional substrate for algal growth (Hoelker et al., 2015; Herren et al., 2017; Phillips et al., 2019), which could increase demand for certain nutrients within the sediment. Furthermore, a laboratory experiment from our group demonstrated negative density-dependence in survival and growth of Tanytarsini larvae under light- and phosphorous-limiting conditions (Wetzel et al., 2021). Therefore, it is plausible that high larval densities at the source sediment locations were associated with high production by benthic algae, which in turn could have resulted in the depletion of certain nutrients, thereby reducing primary production and subsequent emergence for the next generation (i.e., the generation that was the focus of our experiment). Unfortunately, our experimental results do not provide the requisite evidence to test this hypothesis.

The degree of coupling between primary and secondary production plays a central role in governing material and energy flows through aquatic ecosystems (Chapin et al., 2006; Rüegg et al., 2021). Therefore, characterizing the processes that either promote or inhibit such coupling is a topic of general interest. Ecosystems with high primary productivity, such as Mývatn (McCormick et al., 2021), are expected to have a strong coupling between primary and secondary production, since the influence of allochthonous inputs and detritus are likely to be small relative to contemporary autochthonous production (Wallace et al., 1997; Marczak et al., 2007; Rosi-Marshall et al., 2016). We identified a coupling between primary and secondary production for Mývatn’s *T. gracilentus* population, although the coupling was weak. Our results suggest that accumulated organic material in the sediment is sufficiently abundant that modest emergences are still possible when fresh production is greatly reduced. These results illustrate that primary production and consumer dynamics can be partially decoupled even in systems with low external inputs.

## Acknowledgments

This work was supported by National Science Foundation grants DEB-1052160, DEB-1556208 to Anthony R. Ives, and Graduate Research Fellowships DGE-1256259. The Mývatn Research Station directed by Árni Einarsson provided logistical and scientific support. We thank Aldo Arellano, Aliza Fassler, Caroline Owens, and Rebbeca Wetzel for assistance with fieldwork and sample processing.

## Data and code availability

Data and code will be made available on FigShare upon acceptance.

## Conflicts of interest

The authors have no conflicts of interest to declare.

## Supplemental materials

**Figure S1:**
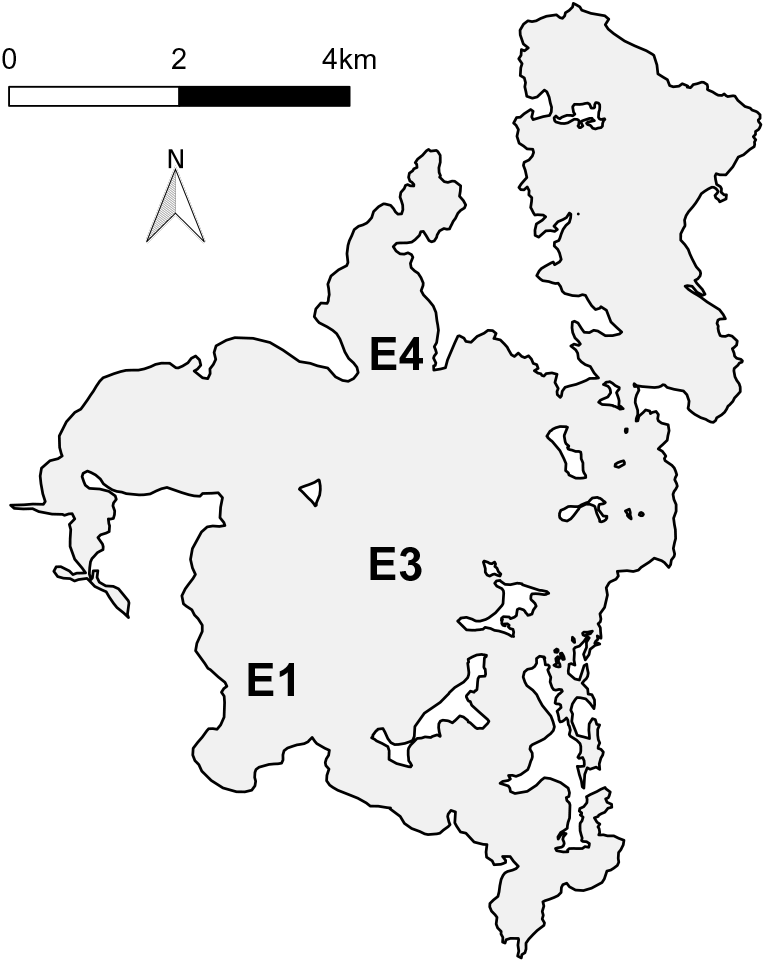
Map of Mývatn showing location of study sites. Gray regions indicate water and white regions indicate land.

**Figure S2:**
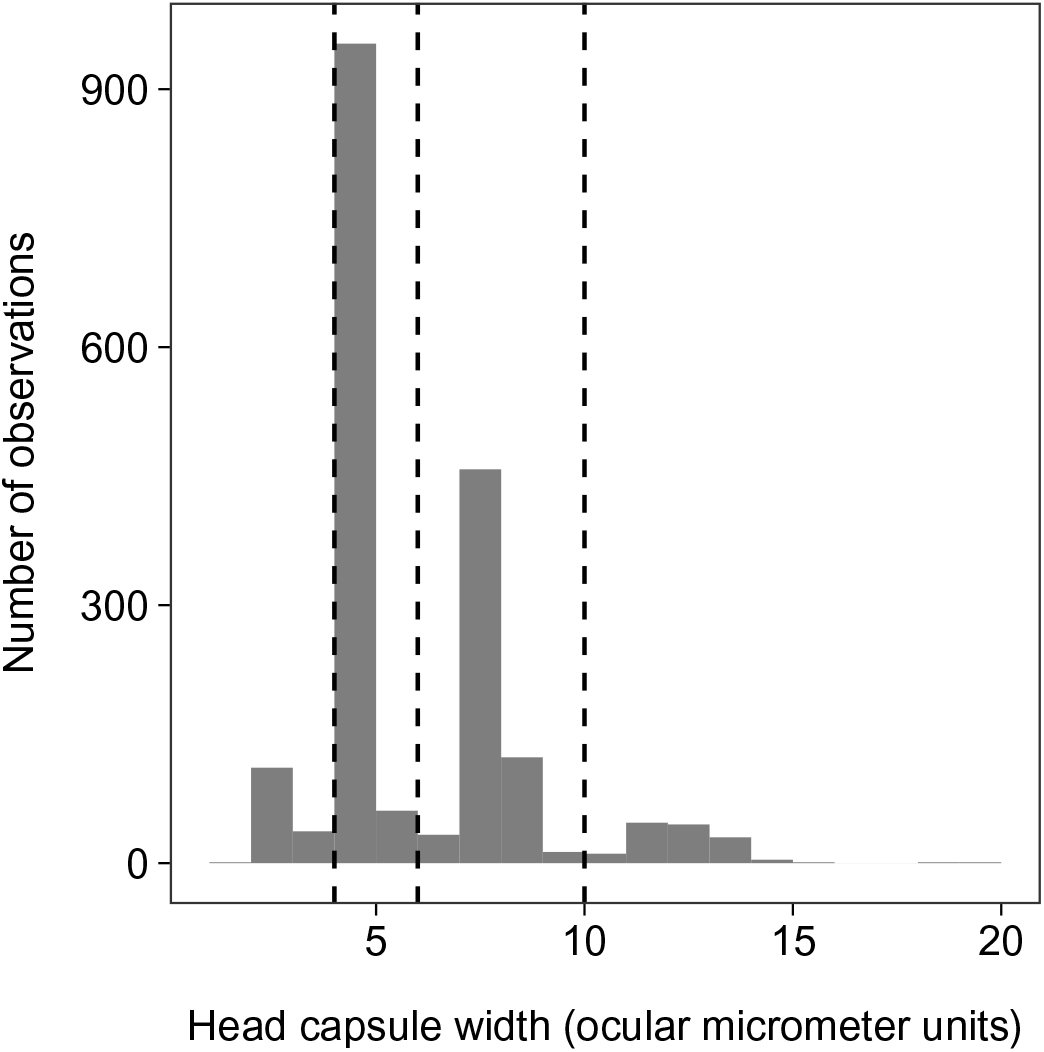
Histogram of larval head capsule width measurements from the experiment. The vertical lines correspond to visually determined threshold for categorizing instars (first: < 4; second: ≥ 4 and < 6; third: ≥ 6 and < 10; fourth ≥ 10).

**Figure S3:**
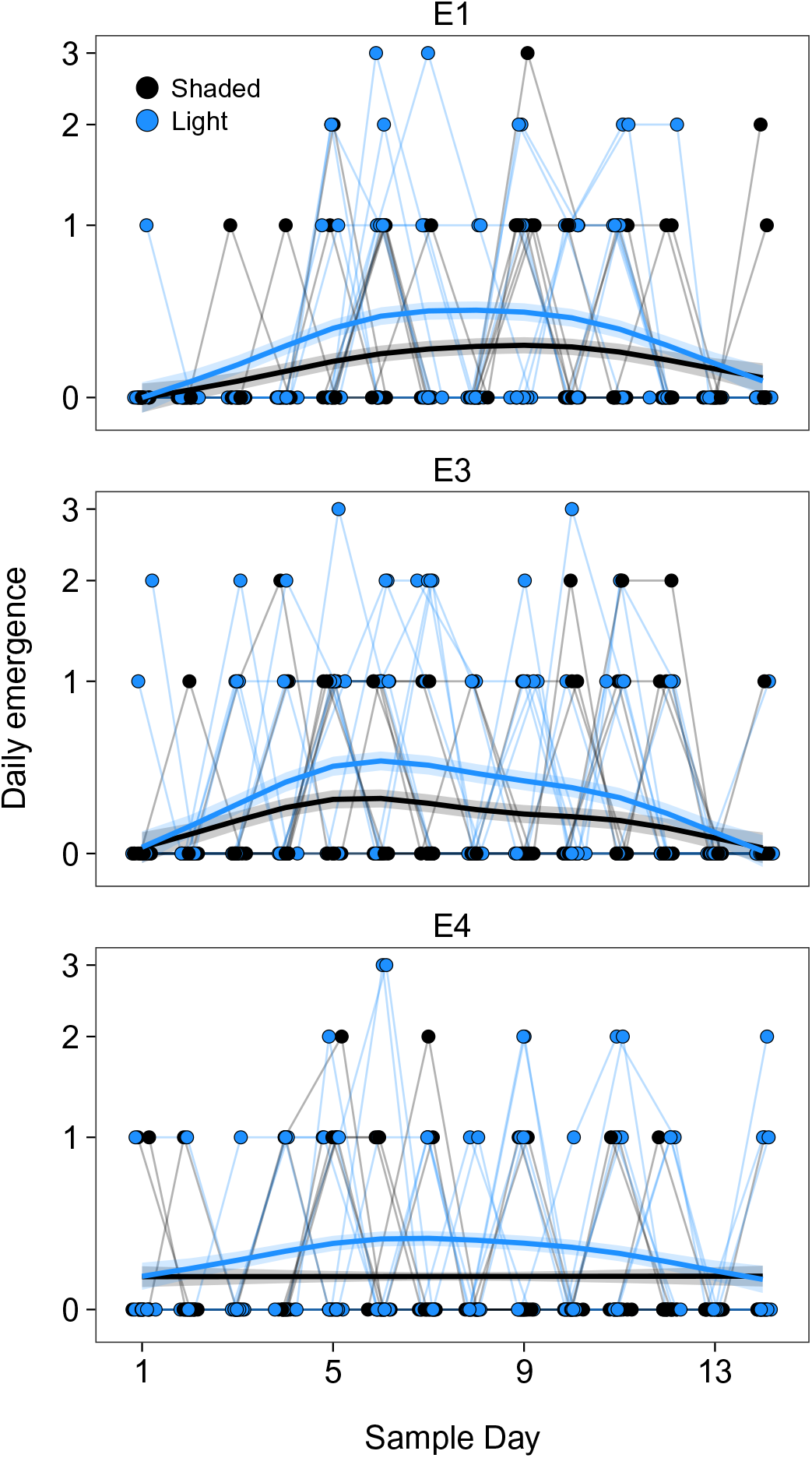
Daily emergence. The lines show fits from a generalized additive mixed model (GAMM) with log(daily count +1) as the response variable and separate spline terms smoothing across days grouped by either sediment source or shading treatment (but not their interaction). We included random intercepts and a lag-1 temporal autoregressive structure for the residuals, grouped by rack (5 levels) and mesocosm (60 levels). The GAMM was fit using the package mgcv in R 4.0.3.

**Table S1:** ANOVA for abundance of Tanytarsini larvae in the mesocosms, with separate models fit for each instar. Counts were log-transformed, with 1 added to all values to accommodate zeros. *P*-values are from *F*-tests with the Kenward-Roger correction.

| Instar | Term | Type III |  | Type II |  |
| --- | --- | --- | --- | --- | --- |
| | | $F_{(ndf,ddf)}$ | $P$ | $F_{(ndf,ddf)}$ | $P$ |
| first | shading | 0.27<sub>(1,50.4)</sub> | 0.608 | 0.85<sub>(1,53.2)</sub> | 0.400 |
|  | site | 0.18<sub>(2,49.2)</sub> | 0.836 | 2.4<sub>(2,51.1)</sub> | 0.098 |
| | shading × site | 1.01<sub>(2,49.5)</sub> | 0.348 | | |
| second | shading | 0.55<sub>(1,51.2)</sub> | 0.462 | 1.8<sub>(1,53.9)</sub> | 0.264 |
|  | site | 2.6<sub>(2,49.4)</sub> | 0.083 | 1.1<sub>(2,51.1)</sub> | 0.354 |
| | shading × site | 1.7<sub>(2,49.8)</sub> | 0.197 | | |
| third | shading | 0.21<sub>(1,50.2)</sub> | 0.645 | 4.0<sub>(1,52.8)</sub> | 0.051 |
|  | site | 2.1<sub>(2,49.2)</sub> | 0.129 | 1.8<sub>(2,51.1)</sub> | 0.179 |
| | shading × site | 2.1<sub>(2,49.41)</sub> | 0.130 | | |
| fourth | shading | 0.003<sub>(1,51.8)</sub> | 0.957 | 4.1<sub>(1,54.5)</sub> | 0.047 |
|  | site | 0.892<sub>(1,49.5)</sub> | 0.416 | 0.27<sub>(2,51.2)</sub> | 0.764 |
| | shading × site | 1.2<sub>(1,50.1)</sub> | 0.298 | | |

**Table S2:** ANOVA for mesocosm GPP. *P*-values are from *F*-tests with the Kenward-Roger correction. These results differ from Table 1 due to the inclusion a variable indicating the mesh treatment (presence-absence). The mesh was intended to exlude colonization of the mesocosms by midge, but it failed as described in the main text.

| Term | Type III |  | Type II |  |
| --- | --- | --- | --- | --- |
| | $F_{ndf,ddf}$ | *P* | $F_{ndf,ddf}$ | *P* |
| PAR | 21<sub>1,55.2</sub> | <0.001 | 18<sub>1,60.1</sub> | <0.001 |
| site | 1.2<sub>2,60.1</sub> | 0.307 | 13<sub>2,50.8</sub> | <0.001 |
| shading | 0.29<sub>1,57.1</sub> | 0.592 | 0.11<sub>1,51.5</sub> | 0.742 |
| date | 26<sub>1,53.7</sub> | <0.001 | 50<sub>1,60.2</sub> | <0.001 |
| mesh | 3.3<sub>1,56.1</sub> | 0.0740 | 2.7<sub>1,51.0</sub> | <0.001 |
| site × shading | 0.73<sub>2,46.3</sub> | 0.489 |  |  |
| site × date | 4.0<sub>2,53.0</sub> | 0.0245 |  |  |
| site × mesh | 0.20<sub>2,46.4</sub> | 0.816 |  |  |
| shading × date | 0.019<sub>1,52.9</sub> | 0.892 |  |  |
| shading × mesh | 0.0072<sub>1,47.5</sub> | 0.933 |  |  |
| date × mesh | 5.7<sub>1,53.1</sub> | 0.021 |  |  |

**Table S3:** ANOVA for abundance of Tanytarsini larvae in the mesocosms. Counts were log-transformed, with 1 added to all values to accommodate zeros. *P*-values are from *F*-tests with the Kenward-Roger correction. These results differ from Table 2 due to the inclusion a variable indicating the mesh treatment (presence-absence). The mesh was intended to exlude colonization of the mesocosms by midge, but it failed as described in the main text.

| Term | Type III |  | Type II |  |
| --- | --- | --- | --- | --- |
| | $F_{ndf,ddf}$ | *P* | $F_{ndf,ddf}$ | *P* |
| site | 13<sub>2,157</sub> | 0.110 | 2.5<sub>2,50.1</sub> | 0.091 |
| shading | 2.2<sub>1,159</sub> | 0.641 | 2.3<sub>1,52.4</sub> | 0.134 |
| instar | 26<sub>3,141</sub> | <0.001 | 144<sub>3,174</sub> | <0.001 |
| mesh | 0.0<sub>3,159</sub> | 0.996 | 0.22<sub>3,50.6</sub> | 0.644 |
| site × shading | 0.63<sub>2,158</sub> | 0.535 |  |  |
| site × instar | 1.1<sub>6,141</sub> | 0.352 |  |  |
| shading × instar | 0.62<sub>3,141</sub> | 0.600 |  |  |
| site × mesh | 2.7<sub>6,159</sub> | 0.074 |  |  |
| shading × mesh | 0.021<sub>6,160</sub> | 0.885 |  |  |
| instar × mesh | 1.5<sub>6,141</sub> | 0.221 |  |  |
| site × shading × instar | 0.50<sub>6,141</sub> | 0.809 |  |  |
| site × shading × mesh | 2.8<sub>6,159</sub> | 0.069 |  |  |
| site × instar × mesh | 1.5<sub>6,141</sub> | 0.198 |  |  |
| shading × instar × mesh | 0.51<sub>6,141</sub> | 0.679 |  |  |
| site × shading × instar × mesh | 1.6<sub>6,141</sub> | 0.156 |  |  |

**Table S4:** ANOVA for number of *T. gracilentus* adults that emerged the mesocosms. Counts were log-transformed, with 1 added to all values to accommodate zeros. *P*-values are from *F*-tests with the Kenward-Roger correction. These results differ from Table 3 due to the inclusion a variable indicating the mesh treatment (presence-absence). The mesh was intended to exlude colonization of the mesocosms by midge, but it failed as described in the main text.

| Term | Type III |  | Type II |  |
| --- | --- | --- | --- | --- |
| | $F_{ndf,ddf}$ | *P* | $F_{ndf,ddf}$ | *P* |
| site | 1.4<sub>2,46.3</sub> | 0.245 | 2.1<sub>2,51.0</sub> | 0.128 |
| shading | 1.4<sub>1,46.9</sub> | 0.239 | 13<sub>1,51.9</sub> | <0.001 |
| mesh | 7.9<sub>1,47.2</sub> | 0.007 | 3.4<sub>1,51.5</sub> | 0.071 |
| site × shading | 0.03<sub>2,46.6</sub> | 0.968 |  |  |
| site × mesh | 1.3<sub>2,46.6</sub> | 0.279 |  |  |
| shading × mesh | 3.2<sub>2,48.3</sub> | 0.078 |  |  |

**Table S5:** ANOVA for wing lenghts of *T. gracilentus* adults that emerged the mesocosms. *P*-values are from *F*-tests with the Kenward-Roger correction. These results differ from Table 4 due to the inclusion a variable indicating the mesh treatment (presence-absence). The mesh was intended to exlude colonization of the mesocosms by midge, but it failed as described in the main text.

| Term | Type III |  | Type II |  |
| --- | --- | --- | --- | --- |
| | $F_{ndf,ddf}$ | $P$ | $F_{ndf,ddf}$ | $P$ |
| date | 4.2<sub>1,196</sub> | 0.109 | 8.9<sub>1,202</sub> | 0.003 |
| site | 2.4<sub>2,45.4</sub> | 0.278 | 0.26<sub>2,39.8</sub> | 0.799 |
| shading | 0.0012<sub>1,38.2</sub> | 0.941 | 0.48<sub>1,44.5</sub> | 0.493 |
| mesh | 4.4<sub>1,60.8</sub> | 0.941 | 1.9<sub>1,43.8</sub> | 0.172 |
| date × site | 4.0<sub>2,195</sub> | 0.047 |  |  |
| date × shading | 0.82<sub>1,188</sub> | 0.420 |  |  |
| date × mesh | 2.4<sub>1,184</sub> | 0.420 |  |  |
| site × shading | 2.3<sub>2,41.9</sub> | 0.190 |  |  |
| site × mesh | 0.49<sub>2,37.6</sub> | 0.190 |  |  |
| shading × mesh | 1.8<sub>2,50.8</sub> | 0.190 |  |  |

## Notes

### Competing Interest Statement

The authors have declared no competing interest.

